# Temperature impacts SARS-CoV-2 spike fusogenicity and evolution

**DOI:** 10.1101/2023.12.20.572501

**Authors:** Jérémy Dufloo, Rafael Sanjuán

## Abstract

SARS-CoV-2 infects both the upper and lower respiratory tracts, which are characterized by different temperatures (33°C and 37°C, respectively). In addition, fever is a common COVID-19 symptom. SARS-CoV-2 has been shown to replicate more efficiently at low temperatures but the effect of temperature on different viral proteins remains poorly understood. Here, we investigate how temperature affects the SARS-CoV-2 spike function and evolution. We first observed that rising temperature from 33°C to 37°C or 39°C increased spike-mediated cell-cell fusion. We then experimentally evolved a recombinant vesicular stomatitis virus expressing the SARS-CoV-2 spike at these different temperatures. We found that spike-mediated cell-cell fusion was maintained during evolution at 39°C, but was lost in a high proportion of viruses evolved at 33°C or 37°C. Consistently, sequencing of the spikes evolved at 33°C or 37°C revealed the accumulation of mutations around the furin cleavage site, a region that determines cell-cell fusion, whereas this did not occur in spikes evolved at 39°C. Finally, using site-directed mutagenesis, we found that disruption of the furin cleavage site had a temperature-dependent effect on spike-induced cell-cell fusion and viral fitness. Our results suggest that variations in body temperature may affect the activity and diversification of the SARS-CoV-2 spike.

**Importance:** When it infects humans, SARS-CoV-2 is exposed to different temperaures (e.g. replication site, fever…). Temperature has been shown to strongly impact SARS-CoV-2 replication but how it affects the activity and evolution of the spike protein remains poorly understood. Here, we first show that high temperatures increase the SARS-CoV-2 spike fusogenicity. Then, we demonstrate that the evolution of the spike activity and variants depends on temperature. Finally, we show that the functional effect of specific spike mutations is temperature-dependent. Overall, our results suggest that temperature may be a factor influencing the activity and adapatation of the SARS-CoV-2 spike in vivo, which will help understanding viral tropism, pathogenesis, and evolution.

## Introduction

In humans, the severe acute respiratory syndrome 2 virus (SARS-CoV-2) mainly replicates in the upper and lower respiratory tracts. In patients, viral antigens have indeed been detected in the nasal epithelium, trachea, or lungs^1,2^. Moreover, under cell culture conditions, SARS-CoV-2 can infect nasal, bronchial, and lung primary human tissues^3^. Infection can cause a wide range of symptoms, from completely asymptomatic infection to mild respiratory disease and even acute respiratory distress syndrome in the most severe COVID-19 cases^4^.

The human airway is characterized by a temperature gradient from 25°C in the nasal cavity to 33°C in the pharynx and 37°C in the lungs. SARS-CoV-2 has previously been shown to replicate better at 33°C than at 37°C^5,6^. In addition to cough, sore throat or fatigue, one of the most common symptoms associated with SARS-CoV-2 infection is fever. Hyperthermia has previously been shown to decrease SARS-CoV-2 replication in vitro^7^. Temperature plays multiple roles in RNA virus transmission, replication, and antiviral immune responses^8^. Low temperatures have been shown to favor spike interaction with its receptor ACE2^9,10^, but the role of hyperthermia in viral entry is not well characterized.

The SARS-CoV-2 spike is one of the three viral proteins exposed on the surface of viral particles, where it is present as an S1/S2 trimer. The S1 subunit mediates the interaction with its receptor ACE2, and S2 contains the fusion machinery necessary for the fusion of the viral envelope with the target cell membrane. The SARS-CoV-2 spike possesses a multi-basic furin cleavage site (FCS) at the S1/S2 junction. This FCS allows cleavage and pre-activation of the spike in producer cells by furin. Upon interaction with ACE2 in target cells, a second proteolytic cleavage fully activates the spike. Depending on the target cell type, this is mediated either by TMPRSS2 at the plasma membrane or by cathepsins in the endosomes. The spike-ACE2 interaction not only mediates entry of viral particles but can also induce cell-cell fusion when the spike on the surface of an infected cell interacts with ACE2 expressed by a non-infected cell^11,12^. Such spike-mediated syncytia have been observed in COVID-19-deceased patients^13^ and have been suggested to play a role in viral spread, pathogenesis, and immune escape^12^. Spike-mediated syncytia formation depends on the presence of the FCS, as its deletion strongly decreases spike fusogenicity^14,15^. However, the effect of temperature on spike-mediated cell-cell fusion is currently unknown.

Since its emergence in humans in 2019, SARS-CoV-2 has evolved into subsequent lineages characterized by different sets of mutations across the whole viral genome. The spike protein is a highly variable genome region, where some variants have accumulated more than 30 mutations. Spike mutations can alter ACE2 affinity, entry pathway, transmissibility, or confer antibody escape^16^. Spike variants also differ in their ability to induce cell-cell fusion. For example, the Alpha, Beta, Gamma and especially Delta variants were associated with increased syncytia formation compared to the ancestral Wuhan-Hu-1 strain^17^. Conversely, the Omicron variants induce less cell-cell fusion^18–20^. Moreover, spike mutations can alter the cellular and tissue tropism of SARS-CoV-2. For example, whereas ancestral strains mainly used TMPRSS2 to enter cells at the plasma membrane, the Omicron variants preferentially use the cathepsin-dependent endosomal entry pathway^18^. This has been suggested to shift the tropism of Omicron from the lower to the upper respiratory tract. Accordingly, Omicron replicates better than ancestral strains in primary nasal and bronchial tissues at 33°C^3^. Conversely, hyperthermia decreases Omicron replication to a greater extent than Delta^21^. Linking spike function, viral evolution, and temperature is therefore critical to understanding SARS-CoV-2 adaptation and pathogenesis.

Here, we investigate the effect of temperature on SARS-CoV-2 spike fusogenicity and, using experimental evolution, we assess how temperature affects spike sequence diversification. We show that high temperatures increase spike-mediated syncytia formation. Moreover, we find that mutations that disrupt the FCS increase in frequency in spikes evolved at 33°C and 37°C, but not at 39°C, because the effects of FCS inactivation on spike-mediated cell-cell fusion, viral entry, and fitness are temperature-dependent. Taken together, these results suggest that temperature affects spike function and is one of the factors influencing the evolution of SARS-CoV-2.

## Results

### High temperatures increase SARS-CoV-2 spike fusogenicity

To measure the effect of temperature on the fusogenicity of the SARS-CoV-2 spike, we used a previously described GFP complementation cell-cell fusion assay^11^ (**Figure 1A**). HEK293T cells expressing two different parts of GFP were mixed and transfected with the SARS-CoV-2 spike (Wuhan-Hu-1 strain) and ACE2. Cells were incubated at 33°C overnight and then transferred to different temperatures (33°C, 37°C or 39°C) for 3h, 6h or 8h. The interaction between spike and ACE2 led to cell-cell fusion and reconstitution of GFP, which was measured by quantitative fluorescence microscopy (**Figure 1B**). This showed that the kinetics of spike-mediated cell-cell fusion was temperature-dependent (**Figure 1C**). The GFP signal observed after 8 h at 39°C was 2.4-fold higher than at 37°C, and 4.8-fold higher than at 33°C. A general linear model of the effects of temperature, experimental block and time (covariate) on the log GFP signal confirmed that temperature significantly increased spike-mediated membrane fusion over time (*p* < 0.0001).

**Figure 1.**
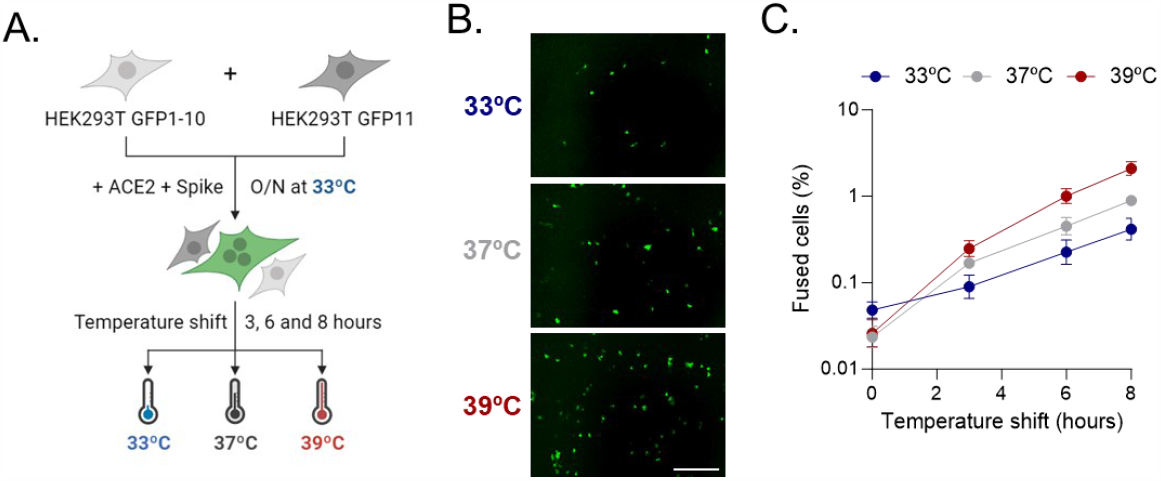
SARS-CoV-2 spike-mediated cell-cell fusion increases with temperature. **A**. Schematic of the cell-cell fusion assay with temperature shift. HEK293T GFP-Split cells were mixed and transfected overnight at 33°C with ACE2 and the SARS-CoV-2 spike. Cells were then shifted to different temperatures (33, 37 and 39°C) for 3, 6 or 8 h and GFP-positive syncytia were quantified. **B**. Representative images of spike-mediated cell-cell fusion after 8 h at different temperatures. Scale bar: 1 mm. **C**. Percentage of spike-mediated cell-cell fusion after shifting transfected GFP-Split cells to different temperatures. Mean and SEM are shown (*n* = 3).

### The evolution of spike fusogenic activity is temperature-dependent

We then investigated whether temperature affected spike diversification using an experimental evolution approach (**Figure 2A**). We obtained a recombinant vesicular stomatitis virus (VSV) modified to express the SARS-CoV-2 spike (strain Wuhan-Hu-1; rVSV-SC2) instead of the VSV G envelope glycoprotein. Using a recombinant VSV virus is a safe, relevant and convenient way to study SARS-CoV-2 spike function and evolution. First, the activity (e.g. entry pathway, recognition by anti-spike antibodies…) of the SARS-CoV-2 spike is similar between a VSV recombinant and a real SARS-CoV-2 virus^22–24^. Second, previous work has already used this system to study the adaptation of the SARS-CoV-2 spike^25^. Finally, this allows to avoid using real SARS-CoV-2 for long-term passaging experiments, which could potentially generate gain-of-function viral variants. After 20 passages in VeroE6-TMPRSS2, we observed that a high proportion of viruses passaged at 33°C and 37°C induced small foci in a titration assay compared to the large foci showed by the founder virus (**Figure 2B**). Large foci consisted of multinucleated syncytia, whereas small foci were indicative of viruses with impaired cell-cell fusion capacity. Such viruses with low cell-cell fusogenicity were observed at low frequency from passage 9-10 at 33°C and 37°C and their proportion gradually increased until passage 15, when their average frequency plateaued at around 60% (**Figure 2C**). In contrast, viruses evolved at 39°C all retained their large-foci phenotype and no fusion-deficient viruses were observed at any passage (**Figure 2C-D**). This demonstrates that temperature affects the phenotypic evolution of the SARS-CoV-2 spike, with hyperthermia (39°C) preventing changes in its cell-cell fusion activity.

**Figure 2.**
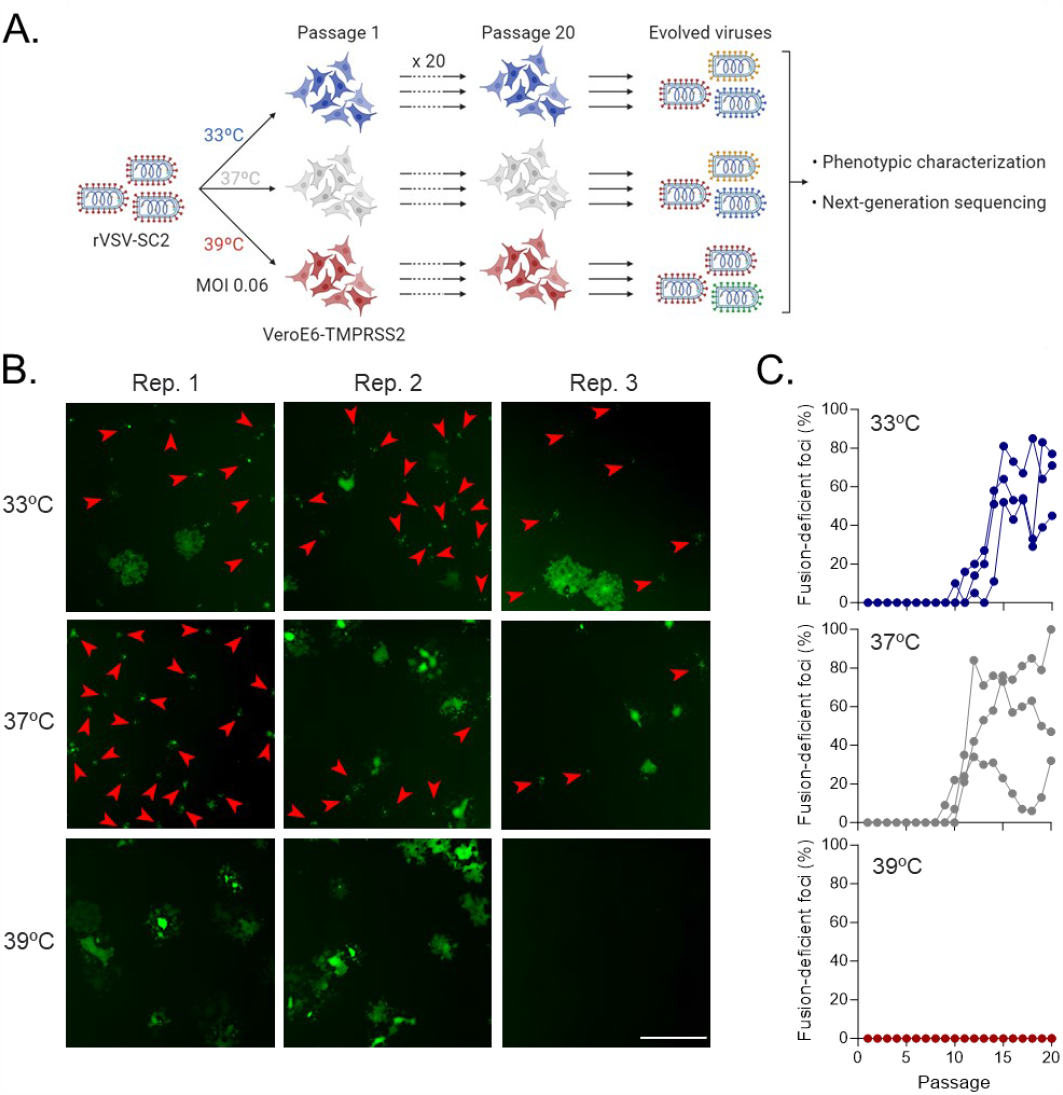
rVSV-SARS-CoV-2 phenotypic evolution is temperature-dependent. **A**. Schematic of the experimental evolution. Recombinant VSV expressing the SARS-CoV-2 spike was passaged 20 times in VeroE6-TMPRSS2 at 33°C, 37°C or 39°C. Three independent evolution lines were performed per temperature. Evolved viral populations were characterized phenotypically and the spike gene was sequenced. **B**. Representative images of the titration of viruses passaged 20 times in VeroE6-TMPRSS2 at different temperatures. Red arrowheads indicate foci of viruses with impaired cell-cell fusion activity. Scale bar: 1mm. **C**. Quantification of the percentage of fusion-deficient foci from the titration of each evolution passage. Each line represents an independent evolution replicate (*n* = 3).

### The evolution of SARS-CoV-2 spike variants is temperature-dependent

To understand the genetic basis of this temperature-dependent phenotypic evolution and to determine whether the spike genetic diversification is affected by temperature, we sequenced by Illumina the evolved viral populations (**Figure 3A**). Most of the sequence variants that arose at a >2% frequency were not commonly observed in nature. An exception was H655Y, present in the Gamma and Omicron variants, which occurred in the three 37°C lineages and one of the 39°C replicates (**Supplementary Table 1**). The T20N mutation, present in the Gamma variant, was also observed at low frequency in all lineages. No high-frequency variant affected the receptor-binding domain (RBD), suggesting that ACE2 affinity is not a strong selective pressure in VeroE6-TMPRSS2. The S2 region was also very rarely affected, with only one high-frequency mutation (L858I) observed in only one lineage (39°C R2). All 39°C lineages had at least one N-terminal domain (NTD) mutation (S50L, W64R and/or K182R) with a frequency >5%. NTD mutations were not observed as often in lineages evolved at 33°C and 37°C, suggesting that hyperthermia may promote NTD diversification.

**Figure 3.**
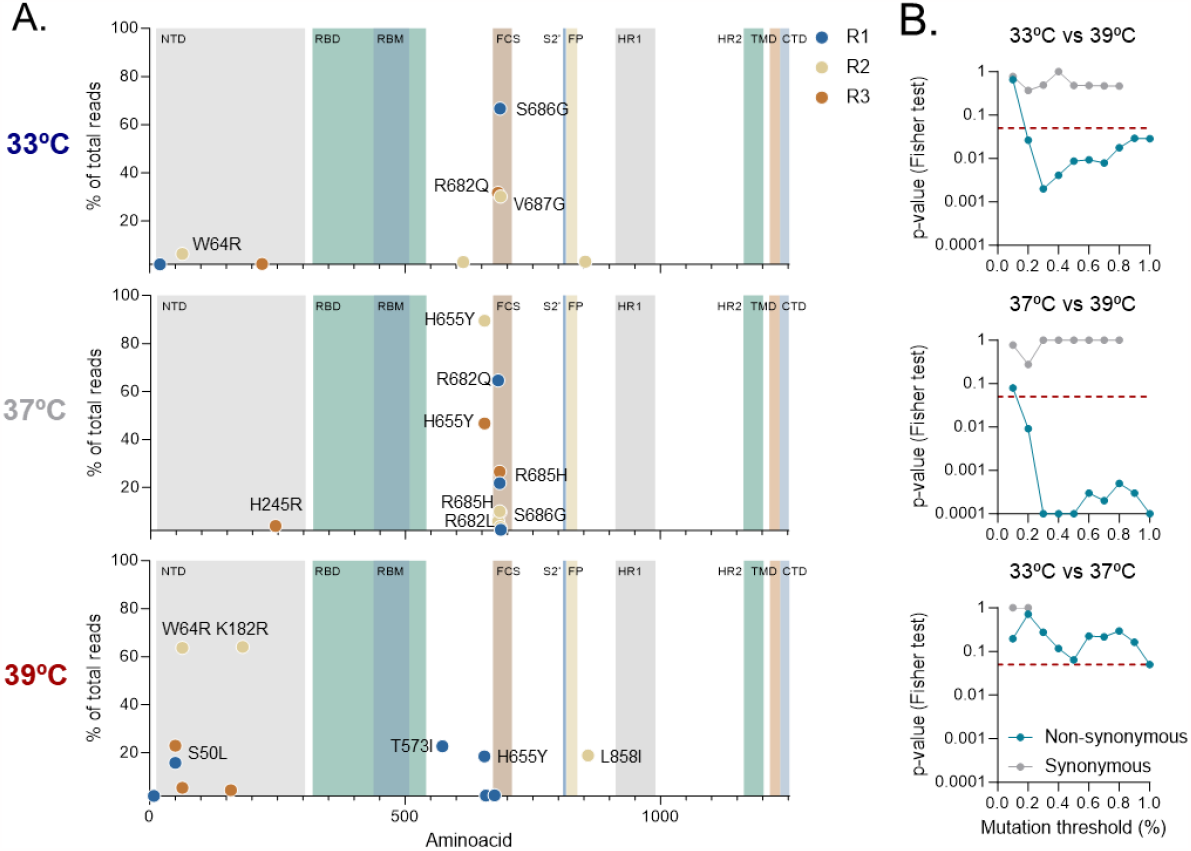
The SARS-CoV-2 spike sequence diversification is temperature-dependent. **A**. Next-generation sequencing of viruses passaged 20 times at different temperatures. Non-synonymous mutations in the spike gene present at a frequency higher than 2% are represented. **B**. The proportion of mutations in the region encompassed by residues 680 to 690 was compared between temperatures using a Fisher test, using different mutation frequency thresholds to include sequence variants (from 0.1 to 1%). This analysis was done separately for non-synonymous (blue) and synonymous (grey) mutations. The red dashed line indicates the statistical significance threshold (*p* = 0.05).

More strikingly, in spikes evolved at 33°C or 37°C, there was a marked clustering of sequence variants in a region encompassed by residues 680 to 690, which contains the FCS. In contrast, this clustering was not observed in spikes evolved at 39°C. Specifically, all 33°C and 37°C lineages had at least one mutation in that region (R682Q, R682L, R685H, S686G and/or V687G) at a frequency >5%, versus only one low-frequency (∽0.1%) mutation (R682L) in the 39°C lineages. To better analyze these differences, we compared the proportion of mutations falling at the FCS region versus the rest of the spike for lineages evolved at different temperatures (**Figure 3B**). Non-synonymous mutations clustered significantly around the FCS in spikes evolved at 33°C or 37°C, but not at 39°C (Fisher’s exact test: *p* < 0.05). This association between the temperature used for evolution and FCS mutation clustering was significant for all variants found at a frequency >0.2% whereas, below this threshold, differences were obscured, probably due to sequencing errors. We also found that the relative frequency of synonymous mutations at the FCS was similar between temperatures, suggesting that the above results were not due to an overall lower genetic diversification at higher temperatures. FCS mutations have previously been described to decrease syncytia formation^14,15,26^ and thus explain the reduced fusogenicity of viruses evolved at 33°C and 37°C. Taken together, this shows that hyperthermia prevents the accumulation of sequence variants around the spike FCS, thereby preserving spike cell-cell fusogenicity, which is otherwise lost during passaging at 33°C or 37°C.

### Temperature-dependent effect of FCS mutations on spike performance

To understand why FCS mutants arose at 33°C and 37°C but not at 39°C, we generated an rVSV-SC2 carrying the S686G mutation, which disrupts the FCS, and measured the effects of this mutation on viral fitness. We confirmed that S686G reduced spike cleavage by furin and spike-mediated cell-cell fusion, in agreement with previous findings^26^ (**Figure 4A-B**). This mutant thus mimics the phenotype observed in the experimental evolution and confirms that S686G is responsible, at least in part, for the reduced fusogenicity of the lineage in which it emerged (33°C R1).

**Figure 4.**
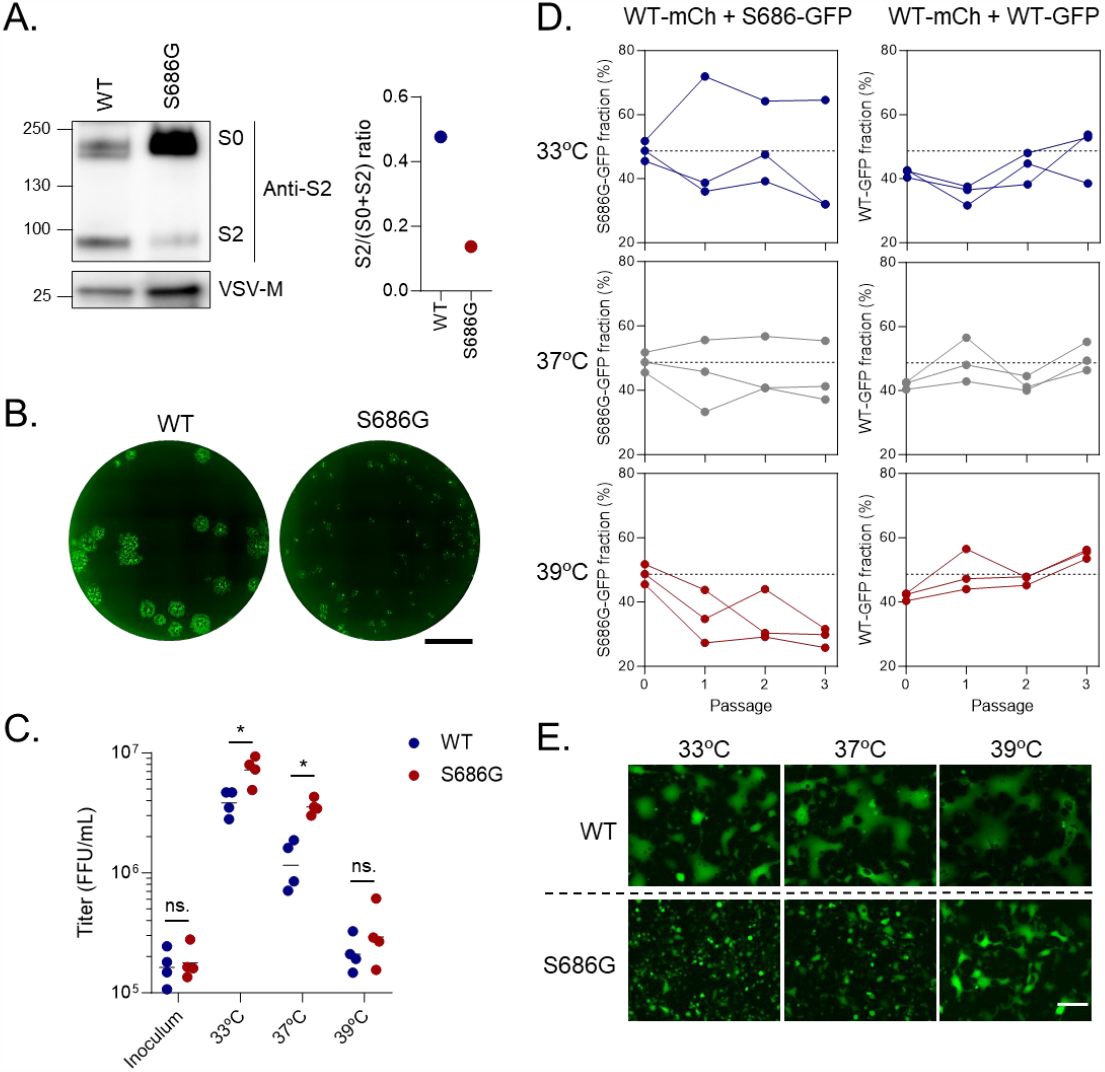
The effect of FCS mutations of spike-mediated cell-cell fusion and viral fitness is temperature-dependent. **A**. Anti-spike S2 western blot of rVSV-SC2 wild-type (WT) and S686G (left panel) and quantification of spike cleavage (right panel). **B**. Foci morphology of rVSV-SC2 WT and S686G mutant. Scale bar: 3.4 mm. **C**. VeroE6-TMPRSS2 were infected by WT or S686G rVSV-SC2 at different temperatures and supernatants were harvested and titrated at 24 hpi. Each dot represents an independent experiment (*n* = 4), and the line indicates the mean. \**p* < 0.05; *ns*. not significant (paired log t-test). **D**. Competition assay between rVSV-SC2-mCherry WT and rVSV-SC2-GFP S686G (left column) or WT (right column). VeroE6-TMPRSS2 were infected with a 1:1 mixture of both virus and three serial passages were performed. The proportion of GFP virus determined at 24 hpi after each passage is represented. Each line represents one independent replicate (*n* = 3). **E**. Representative images of VeroE6-TMPRSS2 infected with rVSV-SC2 WT or S686G for 20 h at different temperatures. Scale bar: 200 μm.

The rVSV-SC2 WT and S686G viruses were then used to infect VeroE6-TMPRSS2 at 33°C, 37°C and 39°C using the same MOI as in the evolution experiment. Fitness was examined in two ways. First, we infected separate wells with each variant and determined the viral titer produced after 24 h (**Figure 4C**). We found that the S686G mutant reached higher titers than the WT at 33°C (2-fold; paired log t-test: *p* = 0.029), 37°C (3.4-fold; paired log t-test: *p* = 0.025), but not at 39°C (paired log t-test: *p* = 0.31). The lack of apparent benefit of this FCS mutation at 39°C may therefore explain why it was not selected at this temperature.

Second, we performed direct competition experiments by infecting each well with a 1:1 mixture of the rVSV-SC2 WT and rVSV-SC2 S686G viruses and propagating these mixed populations for three passages. The two competitors carried different fluorescent reporters in their genomes (mCherry and GFP, respectively), which allowed us to track their frequency by titration and counting of red versus green foci (**Figure 4D**). At 33°C and 37°C, we failed to detect changes in the proportion of each variant after serial transfers (general linear model accounting for experimental block: *p* = 0.324 at 33°C, *p* = 0.174 at 33°C). However, at 39°C, the frequency of the S686G virus gradually decreased with passage number (48.6% in inoculum vs. 29.1% at passage 3), the trend being highly significant (general linear model: *p* < 0.0001). In control assays, we verified that this change in frequency was not due to the different reporters expressed by the WT and S686G viruses (**Figure 4D**). Thus, in competition, the S686G mutation was deleterious at 39°C.

Finally, we found that VeroE6-TMPRSS2 cells infected with the WT rVSV-SC2 fused massively at all temperatures (**Figure 4E**). In contrast, infection with the S686G mutant induced cell-cell fusion in a highly temperature-dependent manner. At 33°C, almost no cell fusion was observed, whereas it was extensive at 39°C and close to that of the WT virus. Cells infected at 37°C showed an intermediate phenotype. We hence hypothesize that the fitness advantage of the S686G mutation at low temperatures may stem from reduced syncytia formation since cell-cell fusion can indeed be detrimental to long-term viral production due to premature cell death^12^. This benefit would be lost at high temperatures, at which FCS disruption fails to prevent syncytia formation efficiently.

## Discussion

We have investigated the effect of temperature on SARS-CoV-2 spike fusogenicity by measuring how it affects spike-mediated syncytia formation. In two different cell types (HEK293T and VeroE6-TMPRSS2), spike-mediated cell-cell fusion increased with temperature. A similar effect of temperature on syncytia formation has been described for HIV-1^27^, Sendai virus27 and the SER paramyxovirus^28^. However, this is not a general rule for syncytia-inducing viruses because cells infected with Varicella Zoster virus fused more at 33°C than at 37°C^29^. The factors that determine the effect of temperature on virus-induced cell-cell fusion remain unclear and warrant further investigation. Temperature increases the fluidity of the plasma membrane, which could facilitate lipid mixing and membrane fusion. However, other processes could be affected by temperature and influence the fusogenicity of the SARS-CoV-2 spike. For example, spike processing is mediated by peptidases (e.g., furin, TMPRSS2, cathepsins) whose activity might be affected by temperature. One study predicted *in silico* that the affinity of the spike/furin interaction may increase with temperature^30^. Temperature could also have a direct effect on spike conformation.

A striking example of the modulation of viral structure by temperature is the transition of dengue virions from a smooth to rugged morphology when the temperature shifts from 28°C to 37°C, affecting the antigenicity and infectivity of viral particles^31,32^. Such a drastic effect of temperature on the structure of the SARS-CoV-2 spike has not been described but low temperatures have been shown to promote the opening of the spike trimer, thereby increasing interactions with ACE2^10,33^.

Syncytia have been observed in lung autopsy specimens from COVID-19-deceased patients^13,34,35^. To our knowledge, syncytia have not been observed in other parts of the respiratory tract. Whether this is because this has not been investigated or because syncytia form specifically in the lung remains unclear. Our data suggest that the higher temperature found in the lower respiratory tract may favor syncytia formation. In addition, COVID-19-associated fever may increase spike-mediated cell-cell fusion. Given the proposed role of syncytia formation in viral pathogenesis (e.g., lung damage, inflammation…), the effect of temperature on syncytia formation deserves further investigation in more relevant primary cell models or in vivo. The latter is complex in humans as histological analyses can only be performed in COVID-19-deceased patients. However, animal models (e.g., non-human primates, hamsters…) infected with SARS-CoV-2 also show syncytia formation^36,37^ and could be used to study the tissue and temperature specificity of SARS-CoV-2 spike-mediated cell-cell fusion.

We used cell-cell fusion to measure spike fusogenicity. However, we did not determine whether high temperatures also increase the entry of SARS-CoV-2 virions. Increased membrane fluidity induced by high temperatures has previously been shown to increase the adsorption and entry of HIV-1 particles^38^. Conversely, high membrane fluidity was detrimental to hepatitis C virus (HCV) entry^39^. Therefore, as with cell-cell fusion, the effect of temperature on viral particle entry cannot be generalized to all viral species and should be further investigated for SARS-CoV-2. The effect of temperature on viral entry may act at different levels. For example, high temperatures inhibit Influenza A virus infection by increasing endosomal pH^40^. Since SARS-CoV-2 has been shown to require an acidic pH to infect cells^41^, this relationship between temperature and pH may be important for SARS-CoV-2 entry. In addition, SARS-CoV-2 variants can use different entry pathways (plasma membrane vs endocytosis), suggesting that their susceptibility to temperature may differ.

It has been shown that FCS disruption is rapidly selected upon passage of SARS-CoV-2 into TMPRSS2-deficient cells^42,43^, but it has been suggested that passaging the virus in TMPRSS2-expressing cells should avoid the emergence of these mutants^44^. We therefore decided to perform our experimental evolution in VeroE6-TMPRSS2 cells to try to avoid this known bias. However, after 20 passages of our rVSV-SC2 in VeroE6-TMPRSS2 at 33°C and 37°C, we repeatedly observed mutations around the FCS region that were associated with a reduced ability to form syncytia. This discrepancy with published results may be multifactorial. First, to our knowledge, no study evaluated the effects of long-term passaging (> 10 passages) of SARS-CoV-2 in VeroE6-TMPRSS2 cells. Second, the levels of TMPRSS2 expressed by our VeroE6-TMPRSS2 may differ from other studies. Finally, most studies describe the adaptation of real SARS-CoV-2 virus, whereas we used a recombinant VSV expressing the SARS-CoV-2 spike, a model that nevertheless captures relevant aspects of SARS-CoV-2 entry^22^. We also note that the spike in our recombinant viruses lacks the last 21 C-terminal amino acids of the S2 subunit, which has been shown to increase viral infectivity and syncytia formation^45^.

The notion that syncytia formation is detrimental to viral fitness in cell cultures is widely supported by the systematic loss of the FCS and spike-induced cell-cell fusion reported here and in previous works^42,43^. Given that spike-mediated cell-cell fusion was more extensive at 39°C, it may be expected that the selective pressure against fusion should also be highest at this temperature. In contrast, the virus did not lose the FCS during evolution at 39°C. The assays performed with the rVSV-SC2 and S686G viruses allowed us to explain these observations since we found that the FCS mutant induced syncytia formation at 39°C, as opposed to 33°C and 37°C. Hence, the fitness advantage of the FCS disruption at 33°C and 37°C was probably lost at 39°C because, at this temperature, cell-cell fusion was induced even by FCS-negative spikes. Whether the virus might find other ways to reduce cell-cell fusion at high temperatures is an open question that could be explored in the future by performing longer evolution experiments.

The fitness assays performed with the rVSV-SC2 WT and S686G viruses provided support to the temperature-dependent effect of FCS mutations on viral fitness. Specifically, the S686G mutation had a positive effect on viral titers at 33°C and 37°C, but not at 39°C. However, the results of the fitness measurements were not fully concordant between mono-infections and direct competition assays in which cultures were coinfected with both variants. The fitness of the S686G virus relative to the rVSV-SC2 WT was lower in coinfection than in mono-infection, such that the mutant was neutral at 33°C and 37°C, but deleterious at 39°C. This discrepancy could be explained in terms of virus-virus interactions. Cell-cell fusion decreases viral yields, but it can also allow rapid spread of the virus in the cell population since the viral release and entry stages of the infection cycle are bypassed. Such rapid spread of the WT syncytia-inducing virus might rapidly exhaust the population of non-infected cells, thus interfering with the growth of the FCS-deficient non-syncytia-inducing virus. Moreover, cells infected with the WT virus could fuse with S686G-infected cells, exerting a negative-dominant effect whereby the S686G mutation would fail to prevent syncytia formation. Consistent with these hypothetical scenarios, despite having a significantly positive effect on viral yield, FCS mutations did not reach fixation in our experimentally evolved populations, which contained a mixture of syncytia-inducing and fusion-deficient viruses. Future research may further address how virus-virus interactions modulate the SARS-CoV-2 spike evolution and, more generally, virally-induced syncytia-induction.

In contrast to the fitness benefit observed in cell cultures, syncytia induction might present some advantages for the virus in vivo, although this is a poorly understood topic. It has been suggested that syncytia formation may allow evasion of antibody-mediated neutralization^46^. Syncytia may also be involved in viral pathogenesis. Indeed, one study showed that spike-mediated syncytia can engulf lymphocytes, inducing their intracellular death and thus representing a possible mechanism underlying COVID-19-associated lymphopenia^47^. It has also been shown that syncytia are prone to premature cell death by apoptosis or pyroptosis^11,48^, which may contribute to the excessive inflammation observed during SARS-CoV-2 infection, but also be detrimental to viral replication, as suggested by our results. It should also be noted that syncytia formation in vivo is likely to be less extensive than in our system since high viral titers, TMPRSS2 expression, and the absence of the E and M proteins facilitate syncytia formation^49^. However, it is interesting to note that the Omicron variants, which rapidly replaced other variants, induce less syncytia formation than the ancestral strains^18–20^. Whether syncytia formation is a selective pressure for SARS-CoV-2 and whether the virus has evolved to avoid the premature cell death associated with extensive cell-cell fusion therefore deserves further investigation.

## Methods

### Cell lines and cell culture

HEK293T-GFP1-10 and HEK293T-GFP11 cells were kindly provided by Olivier Schwartz (Institut Pasteur, Paris, France) and were cultured in the presence of 1 μg/mL of puromycin (Gibco™). VeroE6-TMPRSS2 cells were grown in the presence of 500 μg/mL of G418 (Gibco™). BHK-21 cells were obtained from the ATCC (ATCC CCL-10). BHK-G43 cells were maintained in the presence of hygromycin B (500 μg/mL) and zeocin (1 mg/mL). All cell lines were cultured in DMEM supplemented with 10% fetal bovine serum (FBS), 1% non-essential amino acids, 10 U/mL penicillin, 10 μg/mL streptomycin and 250 ng/mL amphotericin B at 37°C and 5% CO_2_. Cell lines were regularly shown to be free of mycoplasma contamination by PCR.

### Viruses

The plasmid encoding the genome of VSV expressing the SARS-CoV-2 spike (Wuhan-Hu-1 strain; rVSV-SC2) deleted from its last 21 C-terminal amino acids instead of VSV-G (pVSVeGFP-ΔG-Wu-S-ΔCt) was kindly provided by Dr. Ron Geller (CSIC, I2SysBio, Valencia, Spain). The rVSV-SC2 S686G plasmid was obtained through site-directed mutagenesis (Quickchange, Agilent) using a pair of completely overlapping primers (5’-CTCGGCGGGCACGTGGTGTAGCTAGTC-3’ and 5’-GACTAGCTACACCACGTGCCCGCCGAG-3’). Recovery of replication-competent recombinant VSV bearing the SARS-CoV-2 spike (wild-type or S686G) was performed as follows. Briefly, BHK-G43 cells (BHK-21 cells that express the VSV glycoprotein after mifepristone treatment^50^) were seeded at a density of 1.5 x 10^5^ cells/mL in DMEM 5% FBS without antibiotics in 12-well plates (1 mL per well). The following day, the viral genome pVSVeGFP-ΔG-Wu-S-ΔCt was co-transfected with helper plasmids encoding VSV P (25 fmol), N (75 fmol) and L (25 fmol) proteins and the T7 RNA polymerase (50 fmol) using lipofectamine 3000 (Invitrogen™) for 3 h at 37°C. Then the medium was replaced with DMEM 10% FBS supplemented with 10 nM mifepristone to induce VSV-G expression and cells were incubated at 33°C for 36 hours, followed by 48 hours at 37°C. Supernatants from GFP-positive cells were harvested, clarified by centrifugation at 10,000 xg for 10 min, and used to infect a fresh VSV-G-induced BHK-G43 p100 dish culture for amplification. Supernatants were harvested at 24-48 hours post-infection (hpi) and clarified in the same way. To clean viruses from VSV-G, non-G expressing cells were inoculated with the BHK-G43-amplified viruses, inoculum was removed, cells were washed 5 times with PBS and incubated in DMEM containing 2% FBS and 25% of an anti-VSV-G neutralizing monoclonal antibody obtained in-house from a mouse hybridoma cell line. Supernatants were harvested at 24-48 hpi, clarified by centrifugation, and stored at -80°C.

### GFP complementation cell-cell fusion assay

The SARS-CoV-2 HEK293T GFP-split assay was performed as previously described^11^. Briefly, HEK293T-GFP1-10 and HEK293T-GFP11 were mixed at a 1:1 ratio (3 x 10^5^ cells of each cell type per well of a 96-well plate) and transfected with 50 ng of pcDNA3.1-SC2-spike and 50 ng of pCG1-ACE2 plasmid using Lipofectamine 2000 (Invitrogen™) following manufacturer’s instructions. Cells were incubated overnight at 33°C and then shifted at different temperatures (33, 37, or 39°C) for the indicated time (0, 3, 6, or 8h). Images were acquired in an Incucyte^®^ SX5 Live-Cell Analysis System (Sartorius). The percentage of GFP+ area and cell confluence were calculated with the Incucyte^®^ analysis software and the percentage of fusion was calculated as the ratio between GFP confluence and cell confluence.

### rVSV-SC2 experimental evolution

VeroE6-TMPRSS2 cells were plated at a 50% confluence in 6-well plates. The next day, cells were inoculated with 200 μL of virus dilution at an MOI of 0.06. Cells were placed in incubators at 33°C, 37°C or 39°C and agitated every 20 minutes. After 2 hours, 2 mL of DMEM with 2% FBS was added and cells were incubated at different temperatures (33°C, 37°C or 39°C). After 24 hours, supernatants were harvested, cleared by centrifugation (2,000 xg for 10 min), aliquoted, and stored at -80°C. Between each passage, supernatants were titrated as described below. Two passages per week were performed until reaching 20 passages. Three independent replicate evolution lines were performed per temperature.

### Virus titration

VeroE6-TMPRSS2 were seeded in 24-well plates at a 100% confluence for 6-8 hours before being inoculated for 1 hour with serial dilution of virus (100 μL). Cells were then overlaid with 500 μL of DMEM containing 2% FBS and 0.5% agar. After overnight incubation at 37°C, plates were imaged in the Incucyte® SX5 Live-Cell Analysis System (Sartorious). GFP-positive foci were counted manually, and virus titers were calculated as focus forming units (FFU) per mL.

### Next-generation sequencing

RNA from the initial viral stock (P0) and the evolved viruses (passage 20) was extracted using the QIAamp Viral RNA kit (Qiagen) following manufacturer’s instructions. RNA was reverse-transcribed using SuperScript™ IV Reverse Transcriptase (Invitrogen) and a primer recognizing a sequence of the VSV genome upstream of the SARS-CoV-2 spike gene (5’-CTCGAACAACTAATATCCTGTC-3’). SARS-CoV-2 spike gene was amplified by PCR using Phusion Hot Start II DNA polymerase (Thermo Scientific™) and a set of primers recognizing sequences upstream and downstream of the spike gene (Forward: 5’-CTCGAACAACTAATATCCTGTC-3’; reverse: 5’-GTTCTTACTATCCCACATCGAG-3’). PCR products were cleaned using the DNA Clean & Concentrator kit (Zymo Research) and analyzed by Illumina sequencing in an MiSeq machine with paired-end libraries (Novogene). The quality of reads was analyzed with FastQC v0.11.5 (https://www.bioinformatics.babraham.ac.uk/projects/fastqc/). The 15 first and 15 last nucleotides of each read and adapters were removed using Cutadapt (https://cutadapt.readthedocs.io/en/stable/). Reads were then trimmed using the FASTQ quality filter (http://hannonlab.cshl.edu/fastx_toolkit/) and Prinseq-lite 0.20.4 by quality (>Q33), length (>100 nucleotides), and sequencing artifacts (duplications, Ns). The genome of the founder virus was used for mapping and variant calling was performed with FreeBayes^51^.

### Western Blotting

A 1 mL volume of supernatant containing rVSV-SC2 was pelleted by centrifugation at 30,000 xg for 2 hours and lysed in 30 μL of NP-40 lysis buffer (Invitrogen™) for 30 min on ice. Viral lysates were mixed with 4X Laemlli buffer (Bio-Rad) supplemented with 10% β-mercaptoethanol and denaturated at 95°C for 5 min. Proteins were separated by SDS-PAGE on a 4-20% Mini-PROTEAN® TGX™ Gel (Bio-Rad) and transferred onto a 0.45 μm PVDF membrane (Thermo Scientific™). Membranes were blocked for 1 h at room temperature in TBS-T (20 mM tris, 150 mM NaCl, 0.1% Tween-20, pH 7.5) supplemented with 3% Bovine Serum Albumin (BSA; Sigma). Membranes were then incubated for 1 h at RT with the following primary antibodies: mouse anti-SARS-CoV-2 S2 (dilution 1:2,000, clone 1A9, GeneTex) and mouse anti-VSV-M (dilution 1:1,000, clone 23H12, Kerafast). Membranes were washed 3 times with TBS-T and incubated 1h at RT with an HRP-conjugated anti-mouse secondary antibody (dilution 1:50,000, G-21040, Invitrogen). After 3 washes in TBS-T, signal was revealed with SuperSignal™ West Pico PLUS (Thermo Scientific™) following manufacturer’s instructions. Images were acquired on an ImageQuant LAS 500 (GE Healthcare) and analyzed with Fiji software.

### Fitness assays

VeroE6-TMPRSS2 cells were plated at a 50% confluence in 12-well plates. The next day, cells were inoculated with 150 μL of virus dilution at an MOI of 0.06. Cells were incubated at 33°C, 37°C or 39°C and agitated every 20 minutes. After 2 hours, 1 mL of DMEM with 2% FBS was added and cells were incubated at different temperatures (33°C, 37°C or 39°C). After 24 hours, supernatants were harvested, cleared by centrifugation (2,000 xg for 10 min), aliquoted and stored at -80°C until titration as described above.

### Competition assays

VeroE6-TMPRSS2 cells were plated at a 50% confluence in 12-well plates. The next day, cells were inoculated with 100 μL of a 1:1 mixture of rVSV-SC2-mCherry WT and rVSV-SC2-GFP S686G (total MOI = 0.06). A control condition consisting of a 1:1 mixture of rVSV-SC2-mCherry WT and rVSV-SC2-GFP WT (total MOI = 0.06) was included to ensure the observed differences were not due to the expression of GFP vs mCherry. Cells were placed in incubators at 33°C, 37°C or 39°C and agitated every 20 minutes. After 2 hours, 1 mL of DMEM with 2% FBS was added and cells were incubated at different temperatures (33°C, 37°C or 39°C). 24 hours later, supernatants were harvested, cleared by centrifugation (2,000 xg for 10 min), aliquoted and stored at -80°C. Supernatants were then titrated as described above with the difference that both GFP^+^ and mCherry^+^ foci were counted. Supernatants were then used to initiate a new passage by adjusting the total MOI to 0.06. 3 passages were performed in total and the percentage of GFP foci after each passage was quantified. Three independent replicates were performed at each temperature.

### Statistics

Statistics were performed in GraphPad Prism v10 or SPSS v28. All details about statistical tests can be found in the figure legends or in the main text.

## Acknowledgments

We thank all members of the laboratory for helpful discussions about this work. We thank Marc Carrascosa Sàez for help in the generation of the rVSV-SC2 S686G mutant. We thank Iván Andreu-Moreno and Raquel Martínez Recio for technical assistance. This work was financially supported by an ERC Advanced Grant (101019724—EVADER) and a grant from the Spanish Ministerio de Ciencia e Innovación (PID2020-118602RB-I00—ZooVir) to R.S. J.D. is the recipient of an EMBO postdoctoral fellowship (ALTF 140-2021).

## Authors’ contributions

J.D. and R.S. designed research; J.D. performed research; J.D. and R.S. analyzed data; J.D. and R.S. wrote the paper; R.S. provided funding.

